# Using environmental DNA for the detection of *Schistosoma mansoni*: toward improved environmental surveillance of schistosomiasis

**DOI:** 10.1101/530592

**Authors:** Mita Eva Sengupta, Micaela Hellström, Henry Curtis Kariuki, Annette Olsen, Philip Francis Thomsen, Helena Mejer, Eske Willerslev, Mariam Mwanje, Henry Madsen, Thomas Krogsgaard Kristensen, Anna-Sofie Stensgaard, Birgitte Jyding Vennervald

## Abstract

Schistosomiasis is a waterborne, infectious disease with high morbidity and significant economic burdens affecting more than 250 million people globally. Disease control has, with notable success, for decades focused on drug treatment of infected human populations, but a recent paradigm shift now entails moving from control to elimination. To achieve this ambitious goal more sensitive diagnostic tools are needed to monitor progress towards transmission interruption in the environment, especially in low-intensity infection areas. We report on the development of an environmental DNA (eDNA) based tool to efficiently detect DNA traces of the parasite *Schistosoma mansoni* directly in the aquatic environment, where the non-human part of the parasite life cycle occurs. To our knowledge, this is the first report of the successful detection of *S. mansoni* in freshwater samples using aquatic eDNA. True eDNA was detected in as few as 10 cercariae/L water in laboratory experiments. The field applicability of the method was tested at known transmission sites in Kenya, where comparison of schistosome detection by conventional snail surveys (snail collection and cercariae shedding) with eDNA (water samples) showed 71% agreement between the methods. The eDNA method furthermore detected schistosome presence at two additional sites where snail shedding failed, demonstrating a higher sensitivity of eDNA sampling. We conclude that eDNA provides a promising new tool to significantly improve the environmental surveillance of *S. mansoni*. Given the proper method and guideline development, eDNA could become an essential future component of the schistosomiasis control tool box needed to achieve the goal of elimination.

**Significance:** Accurate detection and delineation of schistosomiasis transmission sites will be vital in on-going efforts to control and ultimately eliminate one of the most neglected tropical parasitic diseases affecting more than 250 million people worldwide. Conventional methods to detect parasites in the environment are cumbersome and have low sensitivity. We therefore developed an environmental DNA (eDNA) based method for schistosome detection in aquatic environments. Aquatic eDNA showed higher sensitivity than conventional snail surveys. We conclude that eDNA is a promising non-invasive and sensitive tool for environmental surveillance of schistosomiasis transmission. As the efforts and aims to control the disease are transitioning towards complete transmission interruption, this could be the robust and cost-effective surveillance tool needed in the “end game” of schistosomiasis.

## Introduction

Schistosomiasis is a debilitating infectious disease caused by parasitic worms (blood-flukes) of the genus *Schistosoma* (Fig 1) (1). It is estimated that at least 250 million people globally are infected, and a total of 779 million in 74 countries are at risk of infection (2, 3). With more than 90% of the infected people residing in sub-Saharan African, schistosomiasis is the second most neglected tropical disease (NTD) (4). Over the past decade, the global control strategy has focused on targeted mass drug administration (MDA) programmes leading to reduced worm infections and general improvements in human health (5). However, the focus in schistosomiasis control has now shifted from morbidity control towards transmission-focused interventions (6, 7) as the latest WHO roadmap for disease control aims for elimination (5, 8). This entails a complete interruption of transmission in the environment and thus emphasizes the need for improved environmental surveillance (7). Areas with several years of MDA are expected to have low level parasite transmission, but continued MDAs alone are unlikely to interrupt parasite transmission. Furthermore, as infection levels decrease in the human population with on-going treatment, assessing transmission risk by detecting egg-patent infections in humans becomes less effective (9). Thus, the development and implementation of supplementary environmental surveillance methods to effectively identify presence of schistosome larval stages in the aquatic host snails or directly in aquatic environments is becoming increasingly crucial (7).

As the schistosome parasites critically depend on freshwater snails to complete their lifecycle (Fig 1), environmental surveillance has until now been centered on snail based surveys. This involves collection and identification of host snails followed by light-induced shedding of parasite larval stages from each individual snail (10). Such snail surveys are cumbersome and require substantial specific training and expertise. Furthermore, the sensitivity of this approach is generally low due to only 1-2% of snail populations collected are infected despite high snail abundances and high human infection numbers (11). Even though introduction of DNA techniques for molecular detection of parasite infections in snails recently has revitalized traditional snail monitoring methods (12, 13), extensive snail surveys are still needed to confirm parasite presence. Thus, more sensitive methods to detect schistosome larval stages directly in aquatic environments is still lacking (7, 14).

**Fig. 1.**
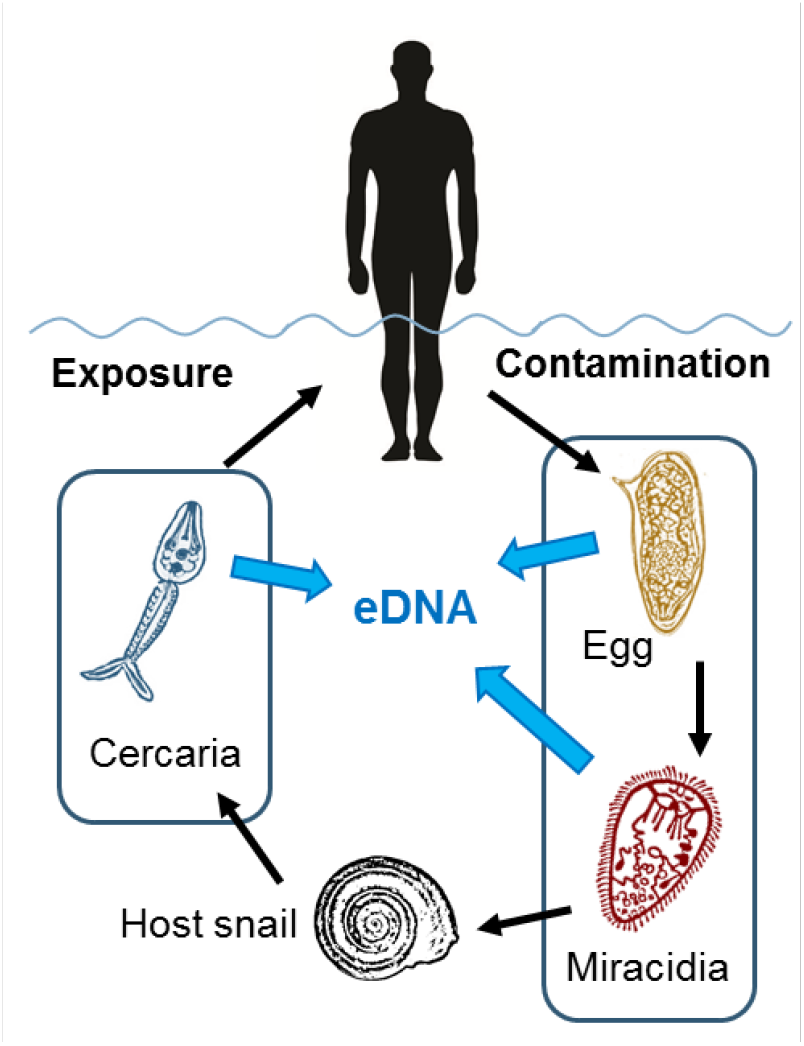
The lifecycle of *Schistosoma mansoni*, illustrating main environmental aspects of transmission related to environmental DNA (eDNA). Infected humans contaminate water sources via feces containing eggs, which hatch into miracidiae larvae infectious to host snails (of the genus *Biomphalaria*). The presence of host snails is essential for further parasite development. After maturation inside the snail, thousands of cercariae are shed into the water, seeking a human host. Each day the emergence, death and decay of the parasite larval stages, and possibly also eggs, contribute to the aquatic pool of eDNA.

To address this challenge, we set out to develop and test a new method to track parasite presence by detecting the traces of DNA, known as environmental DNA (eDNA) (15–17), left behind by the aquatic schistosome larval stages (miracidiae and/or cercariae; Fig 1). Aquatic eDNA in general consists of nuclear or mitochondrial DNA released from organisms via feces, mucous, gametes, skin, hair and carcasses, and can be detected in cellular or extracellular form directly in the environment (18, 19). Our study was inspired by the recent and groundbreaking developments in eDNA research, and its applications in the fields of conservation biology to detect and monitor rare, elusive or invasive aquatic species (20, 21), and the hitherto unexplored potential of eDNA methods in parasitology in general (22).

In the present study, we develop an eDNA-based method to detect the environmental stages of the parasite *Schistosoma mansoni*, causative agent of human intestinal schistosomiasis, in its aquatic environments. We design a species-specific TaqMan qPCR assay, and test this in laboratory tank microcosm experiments to determine assay specificity and sensitivity, as well as schistosome eDNA decay. We then test the applicability and sensitivity of the approach at field sites in Kenya with known history of intestinal schistosomiasis transmission.

## Results

### Species-specific qPCR assay

Primers and probe were designed to specifically target *S. mansoni*, and then successfully validated to be species-specific *in silico* (database blast search, Fig S1), *in vitro* (on tissue-derived DNA extracts of target and non-target schistosome species), and *in situ* (on DNA extracts from tank microcosm and field-collected water samples).

### Tank experiment 1: microcosm and decay of *S. mansoni* eDNA

To validate the species-specificity and sensitivity of the qPCR assay, tank microcosms with varying densities of *Biomphalaria* host snails infected with *S. mansoni* (1, 3 and 6 snails/tank) shedding cercariae into the water was sampled continuously over a 28 day period (Fig 2, Table S1). Schistosome eDNA was detected in water samples at all three snail densities (tank A, B and C in Fig 2) already at the first sampling day and throughout the 28 days, reaching maximum concentration levels of 2.9×10^6^ (1-snail density on day 4), 5.4×10^7^ (3-snail density on day 8), and 2.4×10^7^ (6-snail density on day 8) *S. mansoni* DNA copies/L water (Fig 2). However, a quantitative relationship was not observed between the number of infected snails and the number of schistosome DNA copies detected in the water. To determine schistosome eDNA decay, all snails were removed from the tanks on day 28 and water sampling was continued until day 44 (Fig 2). The parasite eDNA concentrations declined rapidly from concentrations of 1.1×10^6^, 1.5×10^4^ and 3.2×10^6^ DNA copies/L water for snail densities of 1, 3, and 6, respectively, below level of quantification (LOQ; 10 DNA copies/qPCR reaction) and level of detection (LOD; 1 DNA copies/qPCR reaction) (Fig S2). Only the tank with 3 snails significantly fitted the simple exponential decay model and the estimated time for eDNA to degrade below LOQ and LOD (Fig S3) was estimated to be 2.6 and 7.6 days (p<0.05), respectively. All water samples from the two control tanks (D and E in Fig 2) were negative, as well as all day-0 water samples from all tanks (A, B, C, and D in Fig. 2) before addition of snails. All laboratory control samples were also negative leaving no indication of contamination.

**Fig. 2.**
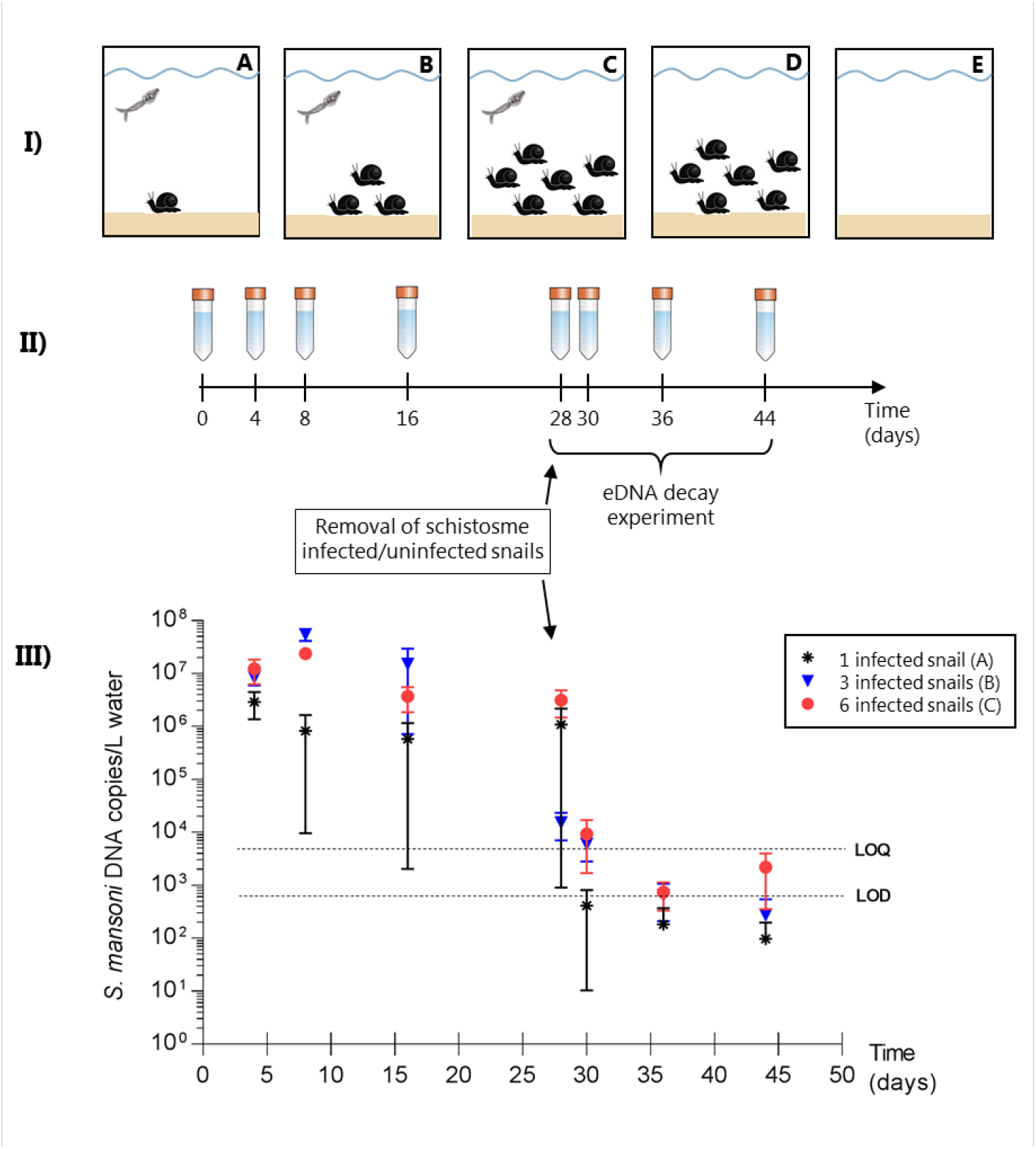
Overview of tank experiment 1 microcosms: **I**) Experimental setup consisted of tanks (*n*=3) of schistosome infected *Biomphalaria* snails at densities of 1 snail (A), 3 snails (B), and 6 snails (C) per tank (4L), and two control tanks with 6 uninfected snails (D) and no snails (E). **II**) Water for eDNA analyses was sampled on day 0 (before adding any snails), 4, 8, 16, 28 (all snails were removed), 30, 36, and 44. From day 28, eDNA decay was measured. **III**) Results showing the concentration of *Schistosoma mansoni* eDNA (copies/L water; ±SEM) on each sampling day with the schistosome infected snail densities of 1, 3 and 6 per tank. The LOQ (10 DNA copy/qPCR reaction) and LOD (1 DNA copy/qPCR reaction) is shown to indicate the position of the data points in relation to these limits (see Fig S1). The two control tanks (D and E) revealed no amplification of *S. mansoni* eDNA and hence not shown in the graph.

### Tank experiment 2: Detection of true eDNA versus whole schistosome cercariae

To determine whether whole cercariae were captured during water sampling, a second tank experiment was performed. Sampling of water with presence of whole cercariae was compared to sampling water with only true eDNA (whole cercariae removed from water) (Fig 3). Results clearly showed that the eDNA method is able to trace true *S. mansoni* eDNA, as opposed to capturing whole cercariae in the water sample (Fig 3). At all three cercariae densities (10, 100 and 1000 cercariae/L water), the removal of cercariae lowered the average level of detected DNA copies considerably. Furthermore, a quantitative relationship was found between the density of cercariae and the amount of schistosome eDNA present in the water.

**Fig 3.**
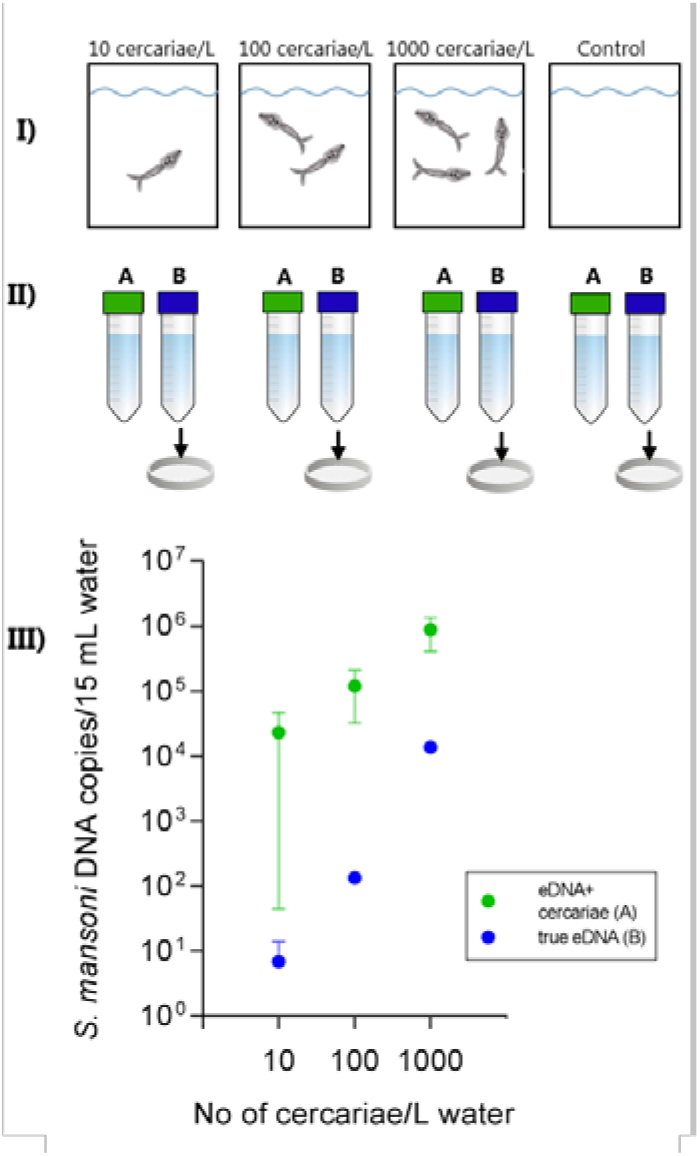
Overview of tank experiment 2. **I**) Experimental tank setup with cercariae density of 10, 100, 1000/L water, and a control tank with water only. **II**) Two series (A and B) of triplicate water samples for eDNA analyses was collected, and the B-series of water samples were filtered to remove whole cercariae. **III**) Results showing the concentration of *Schistosoma mansoni* eDNA (copies/15 mL water; ±SEM) for each cercariae density. The control tank showed no amplification of *S. mansoni* eDNA and hence not shown in the graph.

### Detection of *S. mansoni* at field sites using eDNA and snail surveys

With the eDNA method (qPCR on water samples) *S. mansoni* was detected in water samples from 4/5 sites in Kenya with known ongoing transmission (Fig 4). By comparison, the conventional snail surveys (catching snails and shedding them by means of light-stimulation, followed by PCR) failed to detect schistosome presence at two sites (site 1 and 2) with known transmission (Fig 4; Table 1). At the two sites (site 6 and 7) with no history of transmission, no schistosome eDNA amplified in the water samples and no host snails were found either. Overall the two methods agreed in 71% of the cases (Fig 4; Table S2; Table S3). The overall *S. mansoni* infection rate in the surveyed snail populations in Kenya measured by shedding was 0.4 – 2.2%.

**Fig 4.**
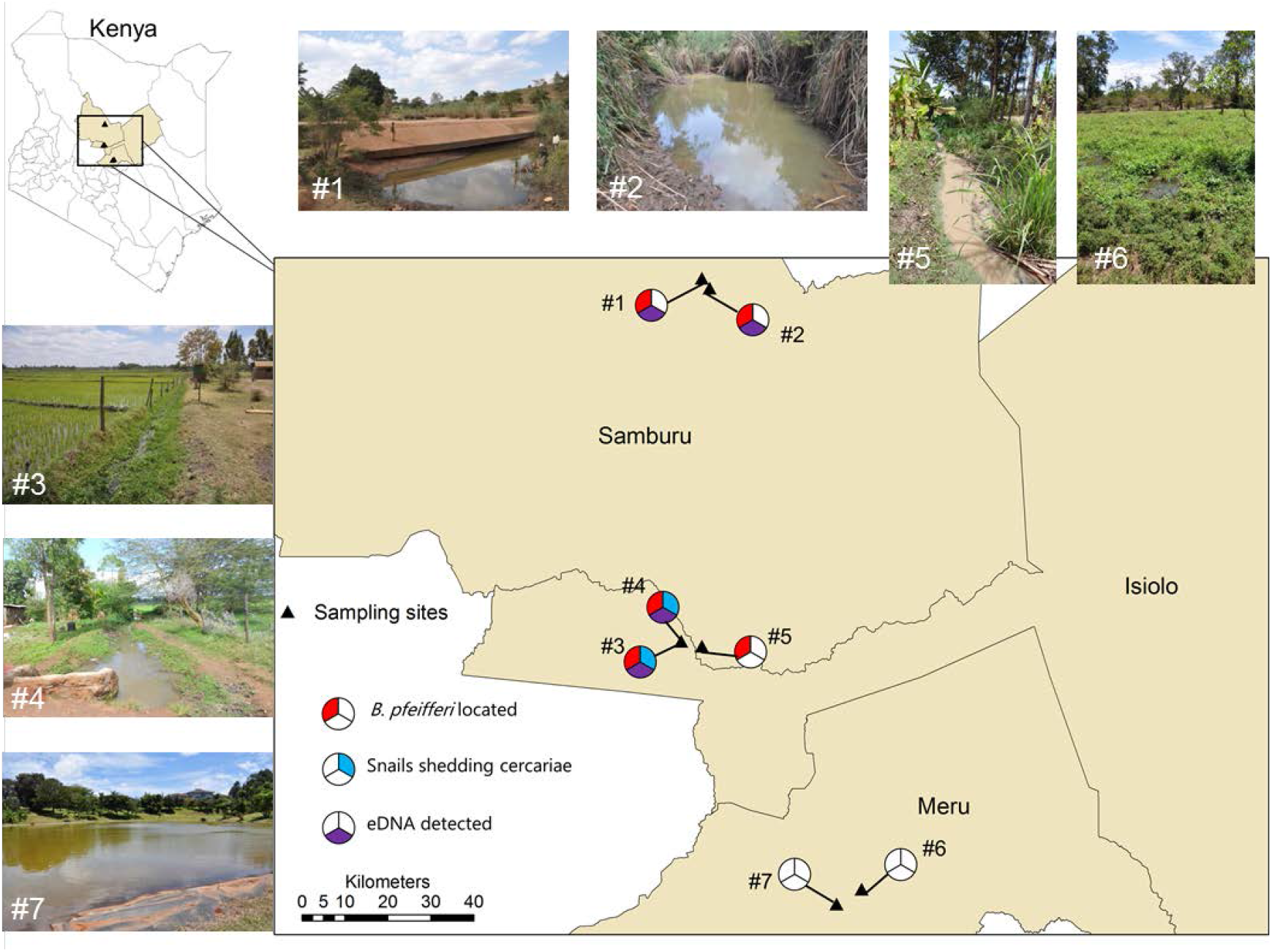
Overview of sampling sites in Kenya and detection results for the conventional snail based survey (*Biomphalariapfeifferi* host snail location and shedding of cercariae) and the eDNA method (water sampling and qPCR analyses). Transmission of *Schistosoma mansoni* is known to be ongoing at site 1-5, whereas site 6-7 has no history of transmission.

Observed naïve detection probabilities at sites where *S. mansoni* was detected was higher for the eDNA method (0.33-0.67) than for conventional snail surveys (0.0004-0.02) (Table 1). The estimated number of water samples required to detect the presence of schistosome eDNA in 95% of the samples at each site ranged from 4-7 samples whereas the conventional snail survey required between 148 and 747 snail specimens of *B. pfeifferi* from each site to achieve a similar level of detection (Table 1).

**Table 1.** Survey metrics and observed naïve detection probabilities of *Schistosoma mansoni* using either snail shedding or eDNA monitoring across seven sites in central Kenya.

| Survey site# | 1 | 2 | 3 | 4 | 5 | 6 | 7 |
| --- | --- | --- | --- | --- | --- | --- | --- |
| Positive samples<sup>a</sup> |  |  |  |  |  |  |  |
| eDNA<sup>b</sup> | 1/3 | 2/3 | 1/3 | 1/3 | 0/3 | 0/3 | 0/3 |
| Snail shedding<sup>c</sup> | 0/11 | 0/2 | 1/240 | 1/45 | 0/73 | 0/0 | 0/0 |
| Detection probability |  |  |  |  |  |  |  |
| eDNA | 0.33 | 0.67 | 0.33 | 0.33 | 0.00 | 0.00 | 0.00 |
| Snail shedding | 0.00 | 0.00 | 0.004 | 0.02 | 0.00 | 0.00 | 0.00 |
| *n* for $P > 0.95^d$ | | | | | | | |
| eDNA | 7 | 3 | 7 | 7 | - | - | - |
| Snails | - | - | 747 | 148 | - | - | - |

<sup>a</sup> Values are the number positive snails or water samples out of the total number of collected samples.
<sup>b</sup> A water sample is designated positive for *S. mansoni* presence if $\geq 1$ qPCR replicate amplified *S. mansoni* DNA.
<sup>c</sup> Number of snails shedding *S. mansoni* cercariae out of the total number of collected snails at each site.
<sup>d</sup> The number of samples (*n*) (snails or water samples) required for the *S. mansoni* site detection probability (*P*) to exceed 0.95 is calculated using the equation $P = 1 - (1-p)^n$ where *p* is the observed naïve detection probabilities at a given site.

### Model based estimates of eDNA detection probabilities in the field

To avoid overestimating the eDNA detection probability at field sites, which can arise from imperfect detection issues (23, 24), the eDNA data was analyzed using a Bayesian multi-scale occupancy model developed specifically for eDNA studies (25). This approach allows the estimation of eDNA occurrence and detection probabilities in relation to various biotic and abiotic factors that may influence detection probability (Table 2) at three hierarchical levels: Ψ (site level), θ (water sample level) and ρ (qPCR replicate level).

**Table 2.** Measured biotic and abiotic factors that potentially influence field site eDNA detection probability, and thus included as covariates in the eDNA occupancy modeling analysis.

| Model covariates | Hypothesized effects | Reference |
| --- | --- | --- |
| *Biomphalaria* snail presence/absence or density. | Could increase parasite eDNA concentration and improve detection in water samples | Present study |
| Presence of snails shedding cercariae | Could increase parasite eDNA concentration and improve detection in water samples ($\theta$). | Present study |
| Salinity | Higher salinity can result in higher availability probability in the water sample ($\theta$) due to increased DNA stability. | (56) |
| Temperature | Affects the physical and metabolic activity of organisms, faster degradation at higher temperatures and lower availability probability in the water sample ($\theta$). | (57, 58) |
| pH | Higher pH has been linked to greater detectability, concentration, and persistence of eDNA in more alkaline waters. | (36, 41, 58, 59) |
| Conductivity | Relates to Total Dissolved Solids (TDS), which can impair eDNA detection due to release of inhibitory substances and their capacity to bind DNA. | (59, 60) |

The occupancy model with the best support (as measured by the posterior predictive loss criterion (PPLC) under squared-error loss and the widely applicable information criterion (WAIC)), included host snail presence as a covariate of eDNA occurrence probability at site level and conductivity as a covariate for eDNA detection probability in qPCR replicate level. A weak positive effect of snail presence on parasite eDNA site occupancy (Bayesian posterior median model estimate 1.85, (95% BCI −0.24; 2.82)), and a negative influence of conductivity on eDNA detection probability in qPCR replicates (posterior median estimate of −0.38 (95% BCI −0.93; 0.15)) was observed (Table 3) However these effects were not significant (95% Bayesian credible intervals for both variables encompassed zero).

Based on the overall model estimated eDNA detection probability at water sample level (θ = 0.35), a total of 7 water samples was estimated to be required to achieve detection probabilities at or above 95% (as calculated using the equation *P* = 1- (1-θ)^n^). The model based estimated number of qPCR replicates required to achieve detection probabilities at or above 95% ranged from 3 to 9 replicates between sites (as calculated using the equation *P* = 1- (1-θ)^n^).

All parameter estimates (posterior medians and 95% credible intervals) for the best fitting eDNA occupancy model can be seen in Table 3. All models and their ranking according to WAIC and PPLC can be seen in Table S4.

**Table 3.** Bayesian posterior estimates of *S. mansoni* eDNA occurrence probability at Kenyan field site (ψ), schistosome eDNA detection probability in a water sample (θ), and schistosome eDNA detection probability in a qPCR replicate (ρ). Parameter estimates (posterior medians and 95% Credible Intervals) are given for each parameter based on for the best fitting eDNA occupancy model (Ψ(snailpres), θ(.), ρ(cond)).

| Site | Snail<br>presence<br>(Y/N) | Conductivity<br>(mS) | $\Psi$ | $\theta$ | $\rho$ |
| --- | --- | --- | --- | --- | --- |
|  |  |  | Median<br>(95% BCI) | Median<br>(95% BCI) | Median<br>(95% BCI) |
| 1 | Y | 1.34 | 0.74 (0.29; 0.97) | 0.35 (0.12; 0.70) | 0.29 (0.08; 0.59) |
| 2 | Y | 0.81 | 0.74 (0.29; 0.97) | 0.35 (0.12; 0.70) | 0.46 (0.28; 0.64) |
| 3 | Y | 0.33 | 0.74 (0.29; 0.97) | 0.35 (0.12; 0.70) | 0.62 (0.34; 0.84) |
| 4 | Y | 0.14 | 0.74 (0.29; 0.97) | 0.35 (0.12; 0.70) | 0.67 (0.33; 0.91) |
| 5 | Y | 0.06 | 0.74 (0.29; 0.97) | 0.35 (0.12; 0.70) | 0.70 (0.33; 0.93) |
| 6 | N | 0.53 | 0.16 (0.01-0.94) | 0.35 (0.12; 0.70) | 0.55 (0.33; 0.75) |
| 7 | N | 0.17 | 0.16 (0.01-0.94) | 0.35 (0.12; 0.70) | 0.67 (0.33; 0.90) |

### Comparison of sampling efforts and associated costs

To investigate the potential cost-effectiveness of the eDNA approach, estimated sampling efforts and associated costs for a further improved eDNA tool and conventional snail sampling was compared for one site. Importantly, a main assumption was that the eDNA method for schistosome detection had been further optimized overcoming the challenges met by the present study. With this in mind, the estimated total effort spent on surveying one site using eDNA was on average approximately half of that using traditional snail collection and shedding (Table 4). The estimated cost for equipment needed for snail surveys and shedding (scoops, trays, beakers) were generally very low and could be re-used several times, whereas enclosed filters and reagents for eDNA analysis cost approximately 165-385 USD per site.

**Table 4.** Estimated efforts (man-hours/site) and costs for materials (USD/site) for sampling and analyses using the eDNA method and the conventional snail based method (snail collection and shedding) to detect schistosomes. Estimations are made based on the lowest and highest number of samples (water samples and snails, respectively) required per site to reach a 95% detection probability (from Table 1).

| Sampling method | eDNA |  | Snail survey<sup>a</sup> |  |
| --- | --- | --- | --- | --- |
|  | 3 water samples/site | 7 water samples/site | 148 host snails/site | 747 host snails/site |
| **Efforts (man-hours):** |  |  |  |  |
| Collection of water samples and filtration/<br>Collection of host snails | 1.0 hrs | 2,3 hrs | 1,3 hrs | 3,3 hrs |
| Lab-work: Extraction to qPCR<sup>b</sup>/Snail shedding | 3.0 hrs | 7.0 hrs | 5,8 hrs | 10.0 hrs |
| **Total effort (per site)** | **4.0 hrs** | **9.3 hrs** | **7.1 hrs** | **13.3 hrs** |
| **Materials (USD):** |  |  |  |  |
| Collection of water samples and filtration | 47 USD | 110 USD | 0 | 0 |
| DNA extraction and qPCR | 118 USD | 275 USD | 0 | 0 |
| **Total cost (per site)** | **165 USD** | **385 USD** | **0** | **0** |

<sup>a</sup> Schistosome infection rate of 2% is assumed for the snail populations.
<sup>b</sup> A total of 9 qPCR replicates for each water sample is assumed, based on the eDNA occupancy model estimates.
*Note:* Here we assume that the required number of snails (148 and 747) is sampled at one site by scooping for 20 min, however this is somewhat an underestimation of man-hours since exploratory sampling at several sites before locating snail populations is often the reality.

## Discussion

To our knowledge, we here present the first successful qPCR-based tool to detect environmental DNA (eDNA) from the snail-borne parasite *Schistosoma mansoni* directly in its freshwater habitat. The demonstrated high level of sensitivity of this eDNA approach to detect schistosome environmental stages will become increasingly important as environmental transmission interruption becomes the measure of true endpoint of schistosomiasis (7).

Earlier attempts to develop a molecular detection method for environmental schistosome stages (26, 27) applied filtering of water using pore sizes appropriate for capturing cercariae, but too large to capture “true” eDNA. In the present study, by employing a state-of-the-art eDNA filtering process we successfully demonstrate that the eDNA method does in fact detect true schistosome eDNA, and not just whole larval stages. This is essential as these stages easily can be missed due to the highly spatial and temporal variation in snail and cercariae density under natural conditions. Moreover, the cercariae are only short-lived with a life expectancy of maximum 24 hours where after they die and degrade beyond detectability if water sample filtering is done with pore sizes too large.

Despite the obvious potential for applying eDNA for environmental surveillance of schistosomiasis, there are a number of limitations and challenges at the current stage that needs discussing. Firstly, for the time being eDNA can only be used to determine the presence (or absence) of schistosomes at field locations, even though knowing the relative densities of parasite infective stages across the infection risk landscape could also be very useful to guide schistosomiasis control efforts. To determine schistosome parasite abundance a quantitative relationship between the number of target organisms and eDNA molecules would be required, as demonstrated in other studies (e.g. 20, 28, 29). In the tank experiment 2, such a relationship between the number of cercariae and concentration of schistosome eDNA was indeed established (Fig 3). However, even though the use of eDNA to quantify species abundances is currently a fast growing field (e.g. 30, 31), some basic issues still remains to be resolved. Importantly, it remains to be resolved how eDNA signals from organism abundance in natural water bodies can be differentiated from organism proximity to where the water samples are taken (32).

Another pressing issue in eDNA studies in general is for how long DNA from an organism is traceable in aquatic environments after removal of the DNA source (33). This is also highly relevant for the applicability of the eDNA method for schistosome detection since the parasite larval stages are relatively short-lived. Our decay experiment (in the tank experiment 1) showed that eDNA detection of cercariae traces in the tank environment would be possible for up to 7 days after the shedding event (Fig. 2). This decay rate is consistent with previous studies estimating the limit of aquatic eDNA detection to be between a couple of days and up to several weeks after removal of the target organism (34–36). However, the decay of schistosome eDNA at actual transmission sites, as compared to controlled tank environments, would probably be faster than a week since the initial DNA concentration of the decay experiment was quite high in comparison to other eDNA decay studies (20, 29). Moreover, increased microbial activity, higher temperatures, and dispersal in natural waters could additionally accelerate the eDNA degradation (32, 37).

Thirdly, under field conditions, it is not possible to determine if the schistosome DNA source originates from cercariae (the human infective stage) or miracidiae (the snail infective stage). This means that the eDNA method cannot at this stage separate detection of contamination (input of miracidiae from infected humans) from exposure potential (snail output of cercariae infective to humans) (Fig 1) (7). Furthermore, we cannot be sure whether the schistosome eDNA arises from living or dead parasite larval stages which also could pose a challenge when assessing real-time transmission (38). However, the latter concern is somewhat unjustified since the short timespan for schistosme eDNA degradation is maximum a week, thus eDNA detection of schistosome presence would indeed represent on-going potential transmission.

A mechanical challenge met during the field testing was filtering of turbid water using pore sizes small enough to capture true eDNA (0.22 μm). Usage of several filter units per water sample (Table S3) was necessary due to clogging of filters even though pre-filtering (with pore size 350 μm) was used to remove larger particles (39). Application of other eDNA capture methods, i.e. other filter types, have not proved to be as efficient as the enclosed filters used in the present field study (40). Additionally, using the enclosed filter units in the field reduces risk of contamination since the filters are never openly exposed. Future studies should focus on how to filter the required volume of water with varying turbidity and simultaneously capture of small DNA fragments in order to keep the number of enclosed filter-units at a minimum, and reduce subsequent laboratory work time.

Lastly, refinement of the eDNA method is needed for use in larger scale schistosomiasis surveillance and control programmes. Thus testing and validation of the method performance under a variety of field conditions and habitats in areas with well-known histories of schistosomiasis transmission is needed. Outstanding questions relate to if and how different habitat types (flowing vs. stagnant waters) or seasonal variation in snail populations and hence also schistosome populations influence schistosome eDNA presence, concentration and detection probability. The next critical step would thus be to develop a panel of sampling guidelines and strategies for eDNA application according to season, habitat type and the type of transmission setting (41) which probably influence the recommended numbers of water samples, the temporal sampling frequency, and the ideal spatial sampling at each habitat type.

Regardless of these challenges, the relatively rapid field collection procedure and simple field equipment combined with a high sensitivity means that eDNA sampling could be widely applicable for broad-scale environmental surveillance of schistosomiasis. The feasibility of eDNA in this context will however depend critically on the associated costs and required efforts of the method. Our results demonstrate that using eDNA to detect *S. mansoni* in the environment may reduce sampling effort and increase detection probabilities relative to the conventional technique (Table 1 and Table 4). In fact, we found that the model estimated “per-sample” eDNA detection probabilities were far greater compared to snail sampling and shedding (requiring as much as 747 snails to be collected at one site). The estimated total sampling efforts for a further optimized eDNA sampling and filtering procedure (Table 4) indicates that man-hours spent per field site is approximately half the time spent for conventional snail surveys, which would make a significant difference in salary expenses. Thus, despite the fact that numerous water samples collected for eDNA analyses would increase the total expenses of the method, the overall cost-effectiveness still appears to be in favor for the more sensitive eDNA method. This is in line with several other studies that have compared eDNA with conventional monitoring methods, and conclude that eDNA can reduce total survey costs (e.g. 42–44). Still, like many other technical sampling methods the eDNA approach entails high start-up expenses which would potentially prevent its implementation under tight surveillance budgets (45).

In the near future, to be able to proceed towards the end-goal of schistosomiasis elimination, the ongoing transition from infection control to transmission control of schistosomiasis is at a critical point where general guidelines are badly needed (7). Recently, WHO published new guidelines for field application of chemical-based snail control (46), but no standard guidelines exist on how to carry out sensitive environmental surveillance. Naturally, eDNA methods cannot stand alone, but in areas with on-going integrated control of MDA and snail control the eDNA method could provide an additional highly accurate means to evaluate control efforts (38). For instance, the eDNA method could be used for closely monitoring of locations declared free of transmission, but where there is a risk of re-establishment of transmission, e.g. due to presence of non-human reservoir hosts where infection might reside undetected by conventional methods. Additionally, early detection of emerging schistosomiasis outside the normally considered endemic range areas using eDNA could be useful to help prevent the disease from spreading. This could for instance be highly relevant in situations where schistosomiasis is moving into new territories, as seen recently in Corsica in Europe due to substantial human migration from endemic transmission areas (47, 48), or due to climate change making new areas suitable for the establishment of both intermediate host snail species and the parasite (49).

Finally, the possibility for eDNA methods to include detection of additional species from the same water samples, i.e. schistosome host snail species or other co-endemic schistosome species, could make the method a true “game-changer” in schistosomiasis environmental surveillance and control. Alternatively, application of the eDNA metabarcoding approach (39) detecting overall species richness in natural environments using high-throughput sequencing of eDNA could be feasible (30). Especially also since snail control is now again emphasized in the plans to eliminate schistosomiasis (38), eDNA detection of schistosome host snail species offers a promising supplement to the conventional snail surveys to help pinpoint “transmission hotspots” (50). The relative ease with which water samples can be collected means that larger geographical areas could be sampled, i.e. through citizen science programmes (44). Thus, eDNA could potentially boost the currently scarce amount of empirical data on host snail and parasite spatio-temporal distributions. These data would allow improved species distribution and risk models, and hence more detailed ‘real-time’ risk maps of schistosomiasis transmission in both emerging and endemic countries, as well as for predicting future risk scenarios under climate change (51). Even though eDNA methods cannot stand alone, given the proper development of the method and associated guidelines, eDNA could become a supplementary and essential future component of the improved environmental surveillance tool needed in the “end game” of schistosomiasis.

## Materials and methods

### Design and validation of species-specific primers

Species-specific primers and probe targeting a 86bp-long sequence in the mitochondrial gene cytochrome oxidase I (COI) of *S. mansoni* (Schiman_COIF: 5’-ATTTACGGTTGGTGGTGTCA-’3; Schiman_COIR: 5’-GAGCAACAACAA ACCAAGTATCA; Schiman_COIprobe: Fam-GGGGTGGCTTTATCTGCATCTGC-BHQ-1-’3) were designed for this study by visual comparison with aligned sequences of *S. mansoni* and other closely related non-target schistosome species occurring in East Africa obtained from NCBI Genbank (Fig S1). Primer and probe sequence motifs were selected with the least theoretical risk of cross-species amplification with non-target species and validated *in silico*. The primer/probe species-specificity was validated *in vitro* by real-time quantitative PCR (qPCR) of genomic DNA tissue extracts from the target species *S. mansoni*, and tested negative for the closely related non-target species *S. rhodaini, S. haematobium* and *S. bovis*.

### Tank experiment 1: Microcosms and eDNA decay

To assess the efficiency and reliability of this proposed eDNA tool the primer-specificity and sensitivity was firstly validated *in situ* in laboratory based tank experiments (microcosms) housing different densities of intermediate host snails, *Biomphalaria glabrata*, infected with *S. mansoni* (see Fig. 2 for experimental setup). Water samples were collected (for ethanol precipitation) before introduction of infected snails (day 0) and at day 4, 8, 16, and 28. Hereafter, snails were removed and sampling of water was continued on day 30, 36, and 44 to examine degradation of schistosome eDNA. Water samples were analyzed using qPCR to quantify DNA amounts and sequenced to confirm *S. mansoni* eDNA.

### Tank experiment 2: Detection of true eDNA versus whole cercariae

To clarify the possible effect of capturing whole cercariae vs. true eDNA when sampling water for *S. mansoni* detection, the tank experiment 2 with different cercariae densities (10, 100 and 1,000 cercariae/L water) were set up (Fig 3). Two series (A and B) of triplicate water samples were taken from each cercariae density and series B samples were filtered (pore size 12 μm) to remove whole cercariae from the water sample, after which all the samples were precipitated. Quantification of *S. mansoni* DNA copies was determined using qPCR.

### Comparison of eDNA method and snail survey in field sites in Kenya

The eDNA method was validated in September 2015 in central Kenya at a total of 7 field sites with known ongoing transmission or with no history of transmission (Fig. 4; Table 1; Table S2). At each site, a water body with human activity was selected and water samples for eDNA analyses was taken before the conventional snail based survey. For eDNA analyses, triplicate water samples of 1 L were taken from each end and the middle of a pond (site 1, 2, 6, and 7) or a selection of a flowing creek/canal (site 3, 4, and 5). A one-liter container with a pre-filter (pore size 350μm) attached to remove large particles was submerged just below the water surface and filled. The water samples were taken standing by the water body edge reaching out wearing long sterile gloves. All field equipment was sterilized in 10% bleach solution and thoroughly dried between sites. Water samples were placed on ice in a dark container immediately after collection until filtering with enclosed Sterivex-filters (0.22μm) using a vacuum pump. Enclosed filters containing eDNA were preserved with RNAlater and kept at −20°C until DNA extraction, following Spens et al.(40). Amplification of *S. mansoni* DNA was done using qPCR.

Conventional snail surveys were performed at each site by catching snails using a scoop for 20 min covering the selected sampling site (52). All specimens of *Biomphalaria pfeifferi* (the intermediate host snail species in central Kenya) were identified based on shell morphology (53) and set up for shedding of cercariae in small beakers placed in the light (sun or artificial) for at least 4 hours as light stimuli induces shedding (10). When large number of snails were scooped the snails were set up for mass shedding of 10 snails in each beaker. All beakers were then visually inspected under microscope and if the fork-tailed schistosome cercariae were detected the 10 snails were singled out in separate beakers to identify the exact snail shedding cercariae. All the host snails were preserved in ethanol 96%. The *S. mansoni* infection of the positive host snails was confirmed using qPCR.

### eDNA decay

An exponential decay model was fitted to the qPCR data from day 28 (set to t=0, as snails are removed) up to 44 from the microcosm experiment, as this is the relationship expected for molecular decay previously shown by Schnell et al (54). The decay model is the following:

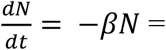

Solving this gives:

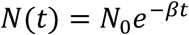

N(t) is the DNA concentration at time t (days). The two parameters N_0_ (initial DNA concentration at t=0) and β (decay constant) were estimated by the nls function in R (vers. 3.4.4), resulting in the values N_0_= 15.19 and β=0.46 for *S. mansoni* in the 3-snail aquaria (tank B in Fig. 2). These parameters were used to calculate after how many days (t) DNA levels would reach beyond level of quantification (LOQ) and level of detection (LOD).

### eDNA occupancy modeling

The R package ‘eDNAoccupancy’ v0.2.0 (25) was used to fit Bayesian, multi-scale occupancy models to estimate schistosome eDNA occurrence and detection probabilities. This approach allowed us to estimate parasite eDNA occurrence and detection probabilities at several hierarchical levels, while also taking the potential effects of environmental covariates into account. The nested survey design in the present study are common for many eDNA surveys (23, 25, 55), and included:

(i) the site occupancy probability (ψ_i_) defined as the probability of schistosome occurrence at site_i_, (ii) the availability probability (θ_ij_) defined as the probability of schistosome eDNA being available for detection in water sample j given that it is present at site _i_, and (iii) the conditional probability of schistosome detection (ρ_ijk_) defined as the probability of schistosome eDNA detection in qPCR replicate k given that it is present in the water sample j and site i.

Several biotic and abiotic factors may potentially affect eDNA detection, persistence, and degradation according to the eDNA literature (Table 2), and therefore we constructed several models to compare the relative importance of these factors. Specifically, we hypothesized that sites with presence of intermediate host snail species, observed shedding or high density of snails, would have a higher site eDNA occupancy probability (ψ), whereas detection probability in water samples (θ) was hypothesized to decrease with increasing salinity, temperature and conductivity. Finally, higher salinity and conductivity were hypothesized to result in inhibition and therefore decrease detection probability at the qPCR level (ρ) (see Table 2 for summary of potential effects of biotic and abiotic factors).

In total 34 models were constructed which included snail-related covariates at site level (Ψ), and a combination of temperature, conductivity and salinity at the water sample level (θ) and conductivity and salinity at the qPCR replicate level (ρ). All models were fitted by running a Markov Chain Monte Carlo (MCMC) algorithm for 11,000 iterations and retaining the last 10,000 for estimating posterior summaries.

Models were ranked (Table S4) according to posterior predictive loss criterion (PPLC) under squared-error loss and the widely applicable information criterion (WAIC). The best model was then fitted by running the MCMC algorithm for 100,000 iterations and retaining the last 50,000 iterations for posterior value estimation. Convergence of the Markov chain used to compute the model estimates was assessed through trace plots of the parameters (25).

Finally, the equations 1 − (1 − θ ^) = 0.95 and 1 − (1 − *p*) = 0.95 (23, 24) were used to determine the number of water samples and qPCR replicates required to achieve detection probabilities at or above 0.95.

### Comparison of sampling efforts and costs

Assuming that the eDNA method for schistosome detection had been further optimized overcoming the challenges met by the present study, the potential cost-effectiveness of the eDNA approach per site was explored. For each of the two monitoring methods, the total effort (measured in man-hours) was estimated based on the lowest and highest number of samples required per site for 95% detection probability (Table 1). For the eDNA method this includes sample collection as well as lab work, and for the conventional snail survey collection and shedding of snails, and visual inspection of cercariae were included.

The cost of materials and reagents for eDNA analysis was estimated, including the cost of extraction, qPCR reagents, and commercial Sanger sequencing. Availability of a qPCR machine and other lab equipment was assumed, and the cost of various plastics, such as pipette tips and tubes, was not included in calculations. Likewise, the cost of snail sampling and shedding gear, such as metal mesh paddle scoops, plastics, and microscopes, was not included in the cost of snail surveys. Costs for travel, subsistence and salaries were not included in these estimates, as they can vary substantially from country to country.

### Ethical statements

Infection of snails for the microcosm experiment was done with *S. mansoni* parasite material recovered from infected mouse-livers delivered by Professor Mike Doenhof, University of Nottingham, UK. During field sampling, collected host snails were not returned to the sites regardless of infection status due to risk of pre-patent infections in the snails.

## Supporting information

Suppl Mat

## Author contributions

Designed research: MES, MH, AO, PFT, EW, TKK, BJV.

Performed research: MES, MH, HCK, AO, HeMe, MM, HeMa.

Contributed new reagents or analytic tools: MH, PFT, EW.

Analyzed data: MES, PFT, HeMa, ASS, BJV.

Wrote the paper: MES, MH, AO, ASS, BJV, with input from all the other co-authors.

## Supporting Information

Supporting Information Appendix (SI Appendix) and Dataset S1 (excel-file).

## Acknowledgments

We are grateful for the financial support received from the Augustinus Foundation and Knud Højgaards Foundation to carry out this study. The technical staff at Kimbimbi Health Centre in Mwea, and Lise-Lotte Christiansen and Rolf Difborg from University of Copenhagen is thanked for their immense help during field sampling. We thank Susanne Kronborg for helping with experimental snail infections and the tank experiments. At Centre for GeoGenetics Tina Brand, Eva Egelyng Sigsgaard and Steen Wilhelm Knudsen are thanked for laboratory assistance. We thank National History Museum London for providing *Schistosoma rhodaini* worm material. EW is grateful to The Danish National Research Foundation, The Lundbeck Foundation, and KU2016 for financial support and thanks St. Johns Collage, Cambridge, for inspiring scientific discussions. ASS is grateful to Knud Højgaards Foundation for supporting the Research Platform for Disease Ecology, Climate and Health and thanks the Danish National Research Foundation for its support of the Center for Macroecology, Evolution and Climate.

## APPENDIX

### Supplementary Information

**This PDF file includes**:

Figures S1 to S3 Tables S1 to S4
Supplementary text (Materials and Methods)
References for SI reference citations

**Other supplementary materials for this manuscript include the following**:

Dataset S1

**Fig S1.**
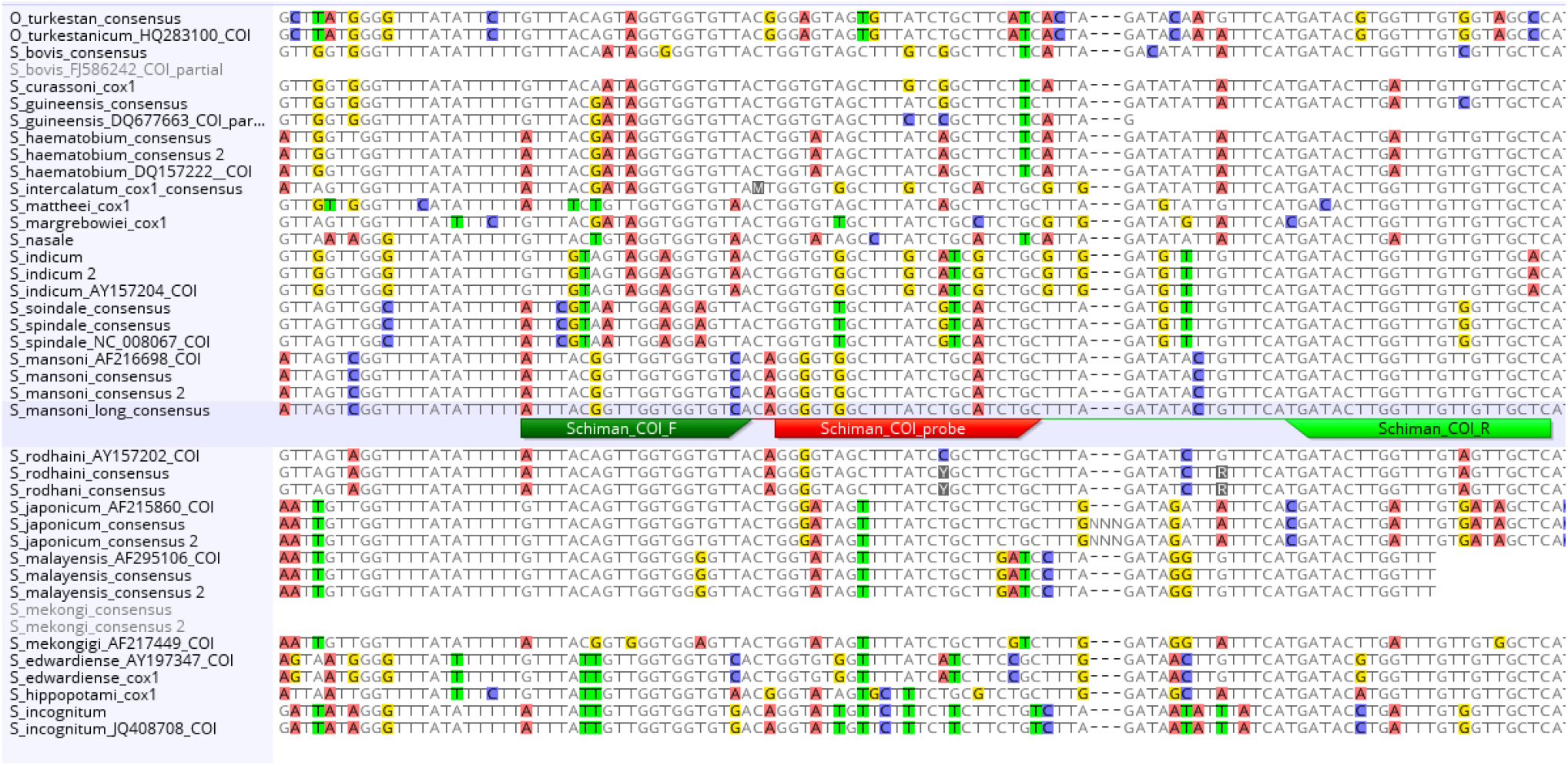
*In silico* assessment of primers and probe targeting *Schistosoma mansoni* showing the number of mismatches in the alignment with other non-target species.

**Fig S2.**
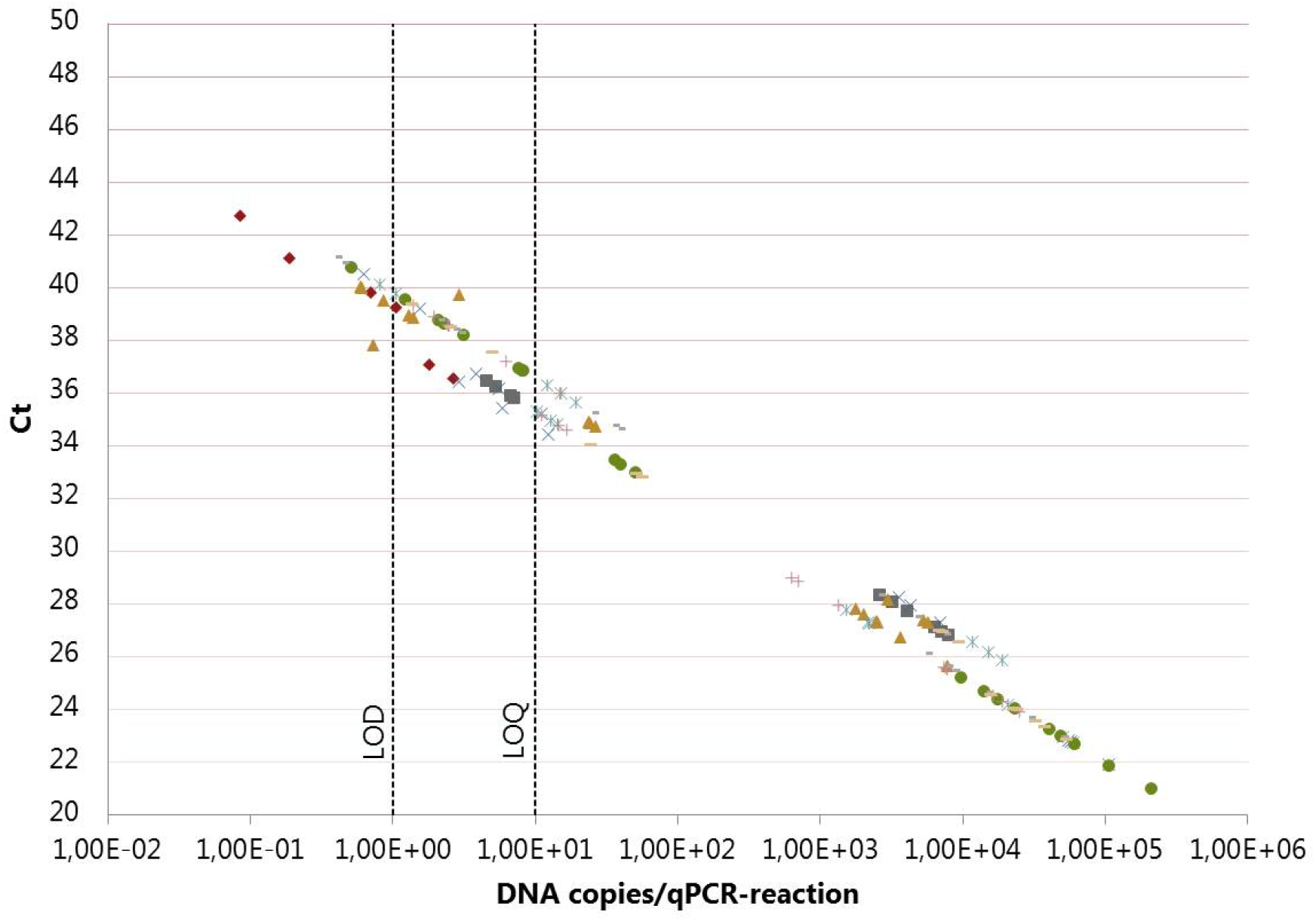
Standard curves for dilution series (10^−1^-10^6^ DNA copies/qPCR reaction) and water samples from tank experiment 1 (microcosm) comparing the concentration of *S. mansoni* eDNA (copies/qPCR reaction) with cycle threshold (Ct). LOQ and LOD were established to be 10 and 1 DNA copy/qPCR reaction, respectively.

**Fig S3.**
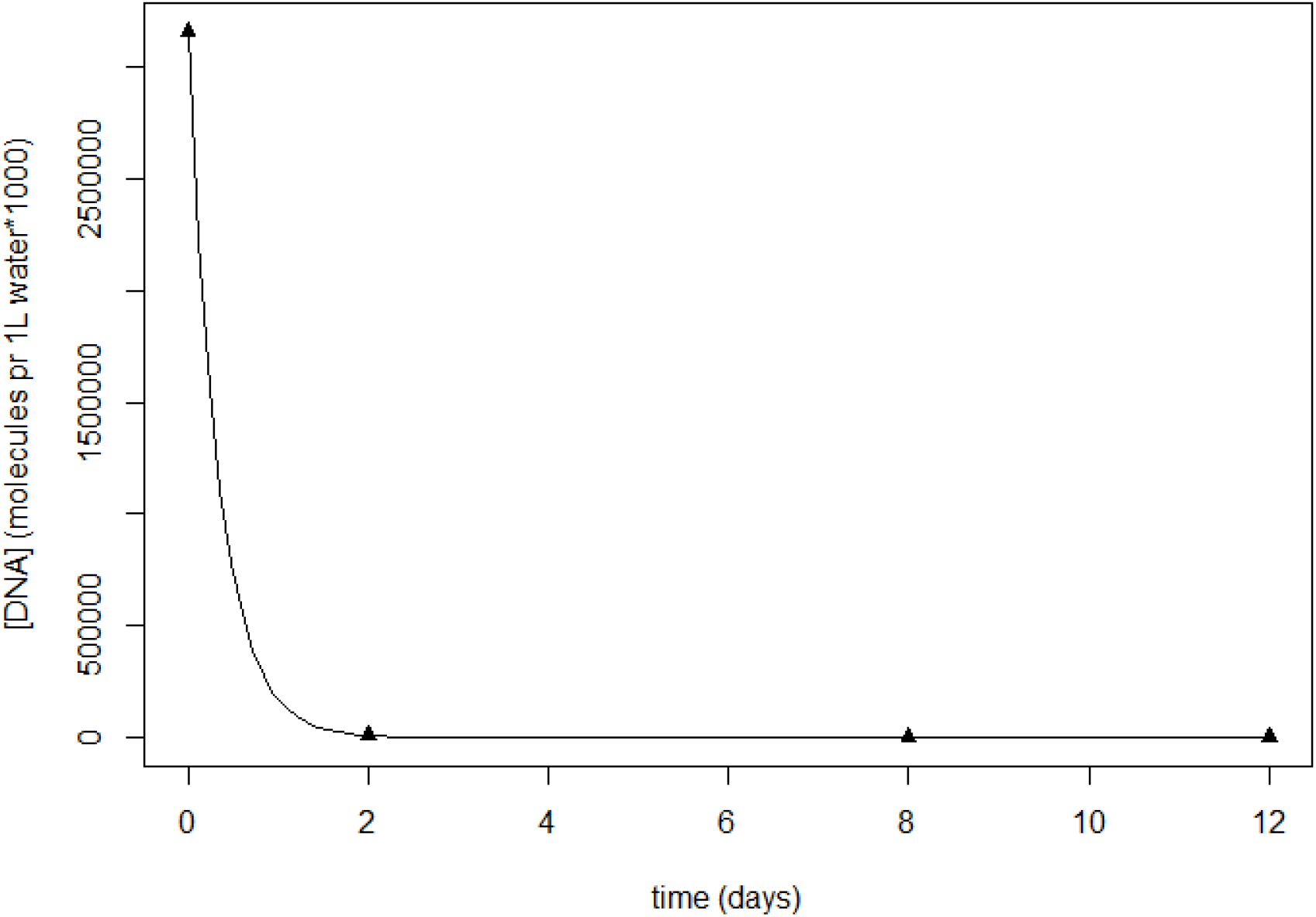
Simple exponential decay model fitted to the *Sschistosoma mansoni* eDNA concentration in tank with density of 3 snails (p<0.05). The aquatic DNA concentration is detected on day 28 in water samples, where after snails were removed. Each data point is the mean of 3 replicate aquaria with 3 qPCR replicates each. Estimated time to decay beyond LOQ is 2.6 days and LOD is 7.6 days. The model fit to the schistosme eDNA concentration in tanks with 1 and 6 infected snails were not significant.

**Table S1:**
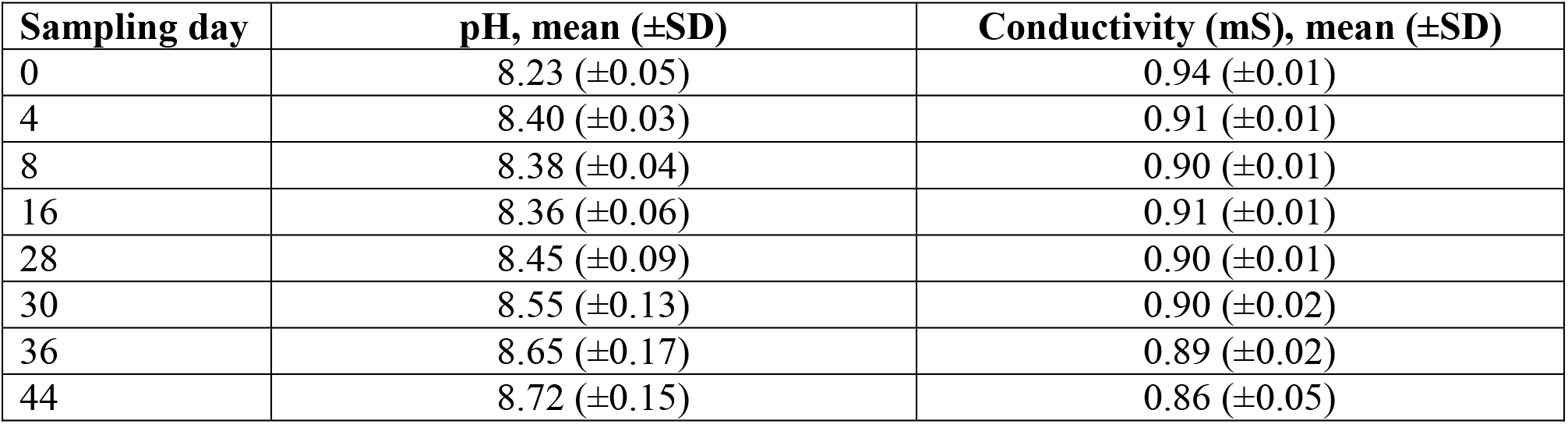
Microcosm water parameters (tank exp. 1). Mean water pH and conductivity values measured on each sampling day during the 44 days of the tank experiment. The values are the total mean (±standard deviation) of 3 replicate tanks for each of the 5 tank types (A, B, C, D and E in Fig 2).

| **Sampling day** | **pH, mean (±SD)** | **Conductivity (mS), mean (±SD)** |
| --- | --- | --- |
| 0 | 8.23 (±0.05) | 0.94 (±0.01) |
| 4 | 8.40 (±0.03) | 0.91 (±0.01) |
| 8 | 8.38 (±0.04) | 0.90 (±0.01) |
| 16 | 8.36 (±0.06) | 0.91 (±0.01) |
| 28 | 8.45 (±0.09) | 0.90 (±0.01) |
| 30 | 8.55 (±0.13) | 0.90 (±0.02) |
| 36 | 8.65 (±0.17) | 0.89 (±0.02) |
| 44 | 8.72 (±0.15) | 0.86 (±0.05) |

**Table S2:** Overview of field sites in Kenya sampled in September 2015, where water sampling for eDNA analyses and conventional snail surveys were compared for detection of *S. mansoni*. Environmental parameters, i.e. water temperature, pH, conductivity and salinity, from each site was used for eDNA occupancy modeling.

| No | Site name | Coordinates | Sampling Date/time | Human water activity | Water flow (m/s) | Water temp. (°C) | pH | Conductivity (mS) | Salinity (ppt) |
| --- | --- | --- | --- | --- | --- | --- | --- | --- | --- |
| 1 | Lower Kalangi River | S01°23.807'<br>E037°22.293' | 14.09.2015<br>3.15 pm | Yes | 0 | 28.6 | 8.17 | 1.34 | 0.67 |
| 2 | Upper Kalangi River | S01°22.571'<br>E037°22.965' | 15.09.2015<br>11.00 am | Yes | 0 | 22.3 | 8.13 | 0.81 | 0.38 |
| 3 | Mianya, canal | S00°38.166'<br>E037°19.408' | 16.09.2015<br>1.30 pm | Yes | 0.3 | 29.4 | 8.15 | 0.33 | 0.16 |
| 4 | Marura Village, canal | S00°38.181'<br>E037°19.416' | 16.09.2015<br>2.15 pm | Yes | 0.17 | 31.3 | 7.98 | 0.14 | 0.07 |
| 5 | Bangladesh, canal | S00°37.542'<br>E037°21.989' | 17.09.2015<br>12.45 pm | Yes | 0.24 | 30.0 | 7.89 | 0.06 | 0.03 |
| 6 | Elephant pond, Meru | N00°07.004'<br>E037°42.054' | 18.09.2015<br>11.45 am | No | 0 | 26.9 | 7.98 | 0.53 | 0.26 |
| 7 | Methodist University pond, Meru | N00°05.130'<br>E037°38.906' | 18.09.2015<br>1.15 pm | No | 0 | 27.8 | 9.80 | 0.17 | 0.08 |

**Table S3.**
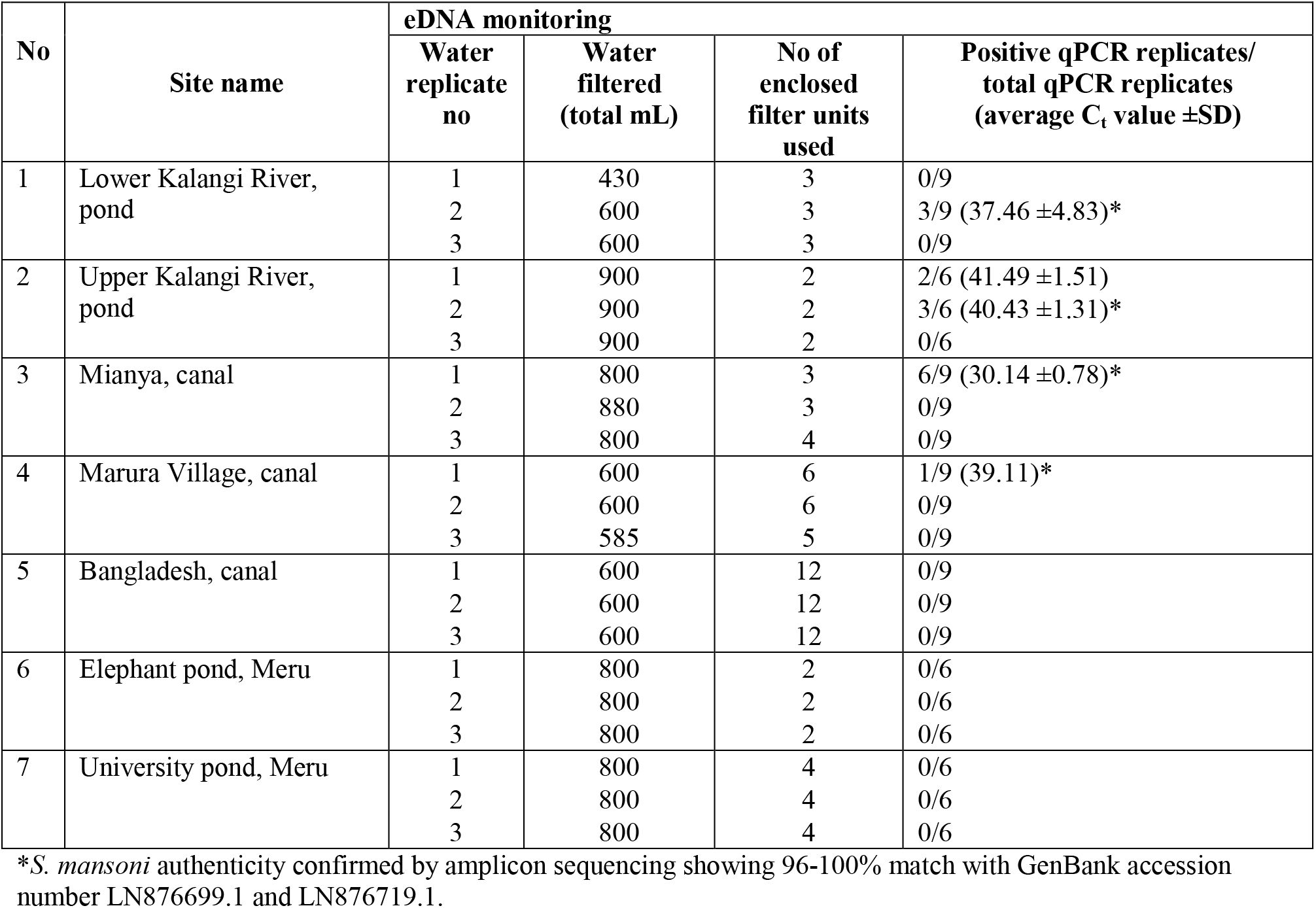
Overview of field water samples collected in Kenya and the following molecular analyses with the species-specific qPCR assay targeting *S. mansoni*. Three replicate water samples were taken at each site and filtered.

**Table S4.** List of 34 fitted eDNA occupancy models and their ranking according to WAIC and PPLC, with the best fitting model highlighted in bold (model number 8).

| model number | model covariates | PPLC | WAIC |
| --- | --- | --- | --- |
| 1 | $(\Psi(.), \theta(.), \rho(.))$ | 15,10 | 0,39 |
| 2 | $(\Psi(\text{snailpres}), \theta(.), \rho(.))$ | 15,00 | 0,38 |
| 3 | $(\Psi(\text{snailpres}), \theta(\text{sal}), \rho(.))$ | 15,60 | 0,41 |
| 4 | $(\Psi(\text{snailpres}), \theta(\text{temp}), \rho(.))$ | 16,00 | 0,45 |
| 5 | $(\Psi(\text{snailpres}), \theta(\text{cond}), \rho(.))$ | 15,40 | 0,41 |
| 6 | $(\Psi(\text{snailpres}), \theta(.), \rho(\text{sal}))$ | 11,80 | 0,37 |
| 7 | $(\Psi(\text{snailpres}), \theta(.), \rho(\text{temp}))$ | 11,70 | 0,38 |
| **8** | **$(\Psi(\text{snailpres}), \theta(.), \rho(\text{cond}))$** | **11,60** | **0,36** |
| 9 | $(\Psi(\text{snailpres}), \theta(\text{sal}), \rho(\text{cond}))$ | 11,90 | 0,37 |
| 10 | $(\Psi(\text{snailpres}), \theta(\text{temp}), \rho(\text{cond}))$ | 13,38 | 0,44 |
| 11 | $(\Psi(\text{snailpres}), \theta(\text{temp}), \rho(\text{sal}))$ | 13,81 | 0,45 |
| 12 | $(\Psi(\text{snailpres}), \theta(\text{cond}), \rho(\text{sal}))$ | 13,57 | 0,46 |
| 13 | $(\Psi(\text{snailshed}), \theta(.), \rho(.))$ | 15,02 | 0,38 |
| 14 | $(\Psi(\text{snailshed}), \theta(\text{sal}), \rho(.))$ | 15,33 | 0,40 |
| 15 | $(\Psi(\text{snailshed}), \theta(\text{temp}), \rho(.))$ | 16,09 | 0,45 |
| 16 | $(\Psi(\text{snailshed}), \theta(\text{cond}), \rho(.))$ | 15,26 | 0,40 |
| 17 | $(\Psi(\text{snailshed}), \theta(.), \rho(\text{sal}))$ | 11,90 | 0,37 |
| 18 | $(\Psi(\text{snailshed}), \theta(.), \rho(\text{temp}))$ | 15,60 | 0,43 |
| 19 | $(\Psi(\text{snailshed}), \theta(.), \rho(\text{cond}))$ | 11,80 | 0,36 |
| 20 | $(\Psi(\text{snailshed}), \theta(\text{sal}), \rho(\text{cond}))$ | 11,90 | 0,38 |
| 21 | $(\Psi(\text{snailshed}), \theta(\text{temp}), \rho(\text{cond}))$ | 13,67 | 0,40 |
| 22 | $(\Psi(\text{snailshed}), \theta(\text{temp}), \rho(\text{sal}))$ | 13,81 | 0,41 |
| 23 | $(\Psi(\text{snailshed}), \theta(\text{cond}), \rho(\text{sal}))$ | 13,77 | 0,40 |
| 24 | $(\Psi(\text{snaildens}), \theta(.), \rho(.))$ | 15,06 | 0,38 |
| 25 | $(\Psi(\text{snaildens}), \theta(\text{sal}), \rho(.))$ | 15,61 | 0,41 |
| 26 | $(\Psi(\text{snaildens}), \theta(\text{temp}), \rho(.))$ | 16,14 | 0,45 |
| 27 | $(\Psi(\text{snaildens}), \theta(\text{cond}), \rho(.))$ | 15,30 | 0,40 |
| 28 | $(\Psi(\text{snaildens}), \theta(.), \rho(\text{sal}))$ | 11,90 | 0,38 |
| 29 | $(\Psi(\text{snaildens}), \theta(.), \rho(\text{temp}))$ | 15,41 | 0,42 |
| 30 | $(\Psi(\text{snaildens}), \theta(.), \rho(\text{cond}))$ | 11,80 | 0,37 |
| 31 | $(\Psi(\text{snaildens}), \theta(\text{sal}), \rho(\text{cond}))$ | 13,34 | 0,42 |
| 32 | $(\Psi(\text{snaildens}), \theta(\text{temp}), \rho(\text{cond}))$ | 12,08 | 0,43 |
| 33 | $(\Psi(\text{snaildens}), \theta(\text{temp}), \rho(\text{sal}))$ | 12,43 | 0,43 |
| 34 | $(\Psi(\text{snaildens}), \theta(\text{cond}), \rho(\text{sal}))$ | 13,48 | 0,46 |

### Text S1: Detailed materials and methods

#### Design and validation of species-specific primers

##### Primer and probe design

Species-specific primers and minor groove binding probes targeting the mitochondrial gene cytochrome oxidase I (COI) of *Schistosoma mansoni* were designed and validated *in silico* by visual comparison with aligned sequences of the *Schistosoma* species occurring in Eastern Africa obtained from NCBI Genbank (Fig S1, above). Primer and probe sequence motifs were selected with the least theoretical risk of cross-species amplification with non-target species. Additionally, the actual species-specificity was validated *in vitro* by real-time quantitative PCR (qPCR) of genomic DNA tissue extracts from the target species and tested negative for *S. rhodaini, S. haematobium* and *S. bovis*.

#### Tank experiment 1: Microcosm and eDNA decay

To assess the efficiency and reliability of this proposed eDNA tool the primer-specificity was first validated in laboratory based tank microcosms housing intermediate host snails, *Biomphalaria glabrata*, infected with *S. mansoni*. Eggs of *S. mansoni* (Puerto Rico strain) were harvested from infected mouse livers (supplied by Professor Mike Doenhof, University of Nottingham) following the protocol of Christensen et al. (1), and 5 miracidiae were used to infect each *B. glabrata* snail (Egypt strain). After a prepatent period of 6 weeks at 23°C all the snails shed cercariae and the microcosm experiment was initiated. The tank setup for the microcosm experiment is shown in Fig. 2. Five different types of aquaria (in triplicates) were set up: A) One infected snail; B) Three infected snails; C) Six infected snails; D) Six uninfected snails; and E) no snails. Two weeks prior to introduction of snails, all aquaria were filled with 4 L ‘pond water’ (tap water softened by presence of guppy fish) and sand/gravel bottom, and added *Daphnia* sp. (purchased from Danish pet-shop) to maintain proper water quality essential for snail survival. During the experiment, snails were fed twice a week with fish food (TetraMin flakes, Tetra GmbH, Germany) and oven-dried organic lettuce. Temperatures were kept constant at 23°C and with a diurnal rhythm of 12 hours of daylight and 12 hours of darkness using artificial light and curtains. On each sampling day pH and conductivity was measured to ensure a stable environment in the tanks throughout the study period (Table S1). Water samples (3x 15 mL) were taken (for ethanol precipitation, following Ficetola et al (2)) before introduction of snails (day 0) and at day 4, 8, 16, and 28 after introduction. Hereafter, snails were removed and sampling of water was continued on day 30, 36, and 44 to examine degradation of schistosome eDNA. All samples were kept at −20°C until DNA extraction.

#### Tank experiment 2: Detection of true eDNA vs. whole cercariae in water samples

To clarify the possible effect of capturing whole cercariae vs. true eDNA when sampling water for *S. mansoni* detection, an experiment with different cercariae densities (10, 100 and 1,000 cercariae/L water) were set up (See Fig. 3). Cercariae was produced as described for tank experiment 1, following Christensen et al. (1). The tanks containing living cercariae in water were left over night at 23°C allowing the cercariae to shed cells into the water to produce eDNA. Two series of water samples were taken from each cercariae density and one of each serie were filtered using polycarbonate filters pore size 12 μm (Nucleopore, Osmonic Inc.) to remove whole cercariae from the water sample. Subsequently all the water samples from the two series were precipitated, following Ficetola et al. (2), and stored at −20°C until DNA extraction.

#### Comparison of eDNA method and snail survey in field sites in Kenya

The eDNA method was validated in September 2015 in the counties Samburu, Isiolo and Meru County in central Kenya at field sites with known ongoing transmission (3) or with no history of transmission (Fig. 4; Table S2). At each possible transmission site a water body with human activity was selected and water sampling for eDNA analyses was taken before the conventional snail survey. At each sampling site water temperature, salinity, pH, and conductivity were measured using a hand-held device (Combo tester, Hanna Instruments, Sweden) to be used in the occupancy analyses. Other snail species present at each site were identified based on shell morphology (4).

For eDNA analyses, water samples of 1 L were taken from the beginning, center, and end (3x 1L) of each of the 7 field sites. A one-liter container with a pre-filter (pore size 350μm, metal kitchen sieve) attached to remove large particles was submerged just below the water surface and filled. The water samples were taken standing by the water body edge reaching out wearing sterile gloved hands. All field equipment was sterilized in 10% bleach solution and thoroughly dried between sites. Water samples were placed in a dark container on ice immediately after collection until filtering with enclosed Sterivex-filters (polyethersulfone; 0.22 μm pore size with luer-lock outlet; Merck KGaA) using a portable vacuum pump (type N811KN.18, P_max_ 2.0 bar, KNF Lab, France). Enclosed filters containing eDNA were preserved with RNAlater and kept at −20°C until DNA extraction, following Spens et al. (5).

Conventional snail surveys were performed at each site by catching snails using a scoop for 20 min covering the selected sampling site (6, 7). All the snails collected were identified to species-level based on shell morphology (4). All the host snail specimens (*Biomphalaria pfeifferi*; the intermediate host snail species in central Kenya), was put up for shedding of cercariae in small beakers placed in the light (sun or artificial) for at least 4 hours, as light stimuli induces shedding (7). When large number of snails was scooped the snails were first set up for mass shedding of 10 snails in each beaker, and then singled out in separate beakers to identify the exact snail shedding cercariae. All beakers were visually inspected under microscope and a snail were designated positive for *S. mansoni* infection if the fork-tailed schistosome cercariae were detected (8). All host snails were kept in ethanol 96%, and subsequently the *S. mansoni* infection in the specific snails shedding cercariae was confirmed by PCR.

#### DNA extraction

DNA extraction and post-PCR work were performed in two separate laboratories assigned for the purposes and equipped with positive air pressure and UV-treatment. DNA was extracted from the enclosed Sterivex-filters using DNeasy Blood and Tissue kit (Qiagen, Carlsbad, CA, USA) with modifications, as described by Spens et al. (5). Extraction blanks were included for all DNA extractions and were tested negative in subsequent PCRs.

#### Quantitative PCR (qPCR)

Primer and probe annealing temperature was optimized by gradient PCR. Reaction volume of 20.0 μL, containing 1.0 μL genomic DNA template retracted from *Schistosoma mansoni* tissue, 10.0 μL of 2.0 U/μL TaqMan^®^ Environmental Master Mix 2.0 (Life Technologies), 1.0 μL of 10.0 μM of each primer (forward and reverse), and 7 μL ddH_2_O. Thermal conditions started with an initial preheat at 95°C for 10 minutes, followed by 32 cycles of (95°C for 30 seconds, gradient temperature for 30 seconds, and 72°C for 30 seconds), and a final extension at 72°C for 10 minutes. The gradient temperatures tested were 56.3°C, 57.4°C, 58.5°C, 59.5°C, and 60.4°C.

The optimized qPCR protocol used for *S. mansoni* eDNA detection and quantification is as follows: One reaction of 25 μL contained 3 μL of template DNA, 12.5 μL TaqMan^®^ Environmental Master Mix 2.00 (Life Technologies), 6.5 μL of UV-treated laboratory-grade water, and 1 μL of each primer and probe (10 μM and 2.5 μM, respectively). Thermal settings included 5 min initial denaturation at 50°C and 10 min at 95°C, followed by 50 cycles of (30 s. at 95°C and 1 min at 57°C) with end-point collections of fluorescence at the 57°C step. All qPCRs were performed on a Stratagene Mx3005P (Thermo Fisher Scientific Inc.). Negative controls (NTC) were included for all PCRs and showed no amplification. To check for inhibition we used an internal positive control (TaqMan^®^ Exogenous Internal Positive Control) adding 2.5 μL of Exo IPC Mix and 0.5 μL of Exo IPC template DNA to the mixture in at least 2 replicates of each sample (3.5 μL dd H_2_O was used in the replicates). No samples showed signs of inhibition (no initial amplification of the dye Vic). For each of the water samples in tank-1 and tank-2 experiments 3 independent qPCR replications were performed, and for each field collected water sample 6-9 independent qPCR replicates. A sample was regarded as positive when a sigmoidal amplification curve was detected in one or more qPCR replicate.

For the tank experiment 1 and 2, qPCR standards for *S. mansoni* were prepared as a dilution series (10^−1^-10^6^ DNA copies/reaction) of purified PCR products on tissue-derived DNA with concentration measured on a Qubit 1.0 fluorometer (Thermo Fisher Scientific Inc.) applying the high-sensitivity assay for dsDNA (Life Technologies, Carlsbad, CA, USA), as described by Agersnap et al. (9). Final concentrations in DNA molecules per volume of water sample were calculated from the standards setting the molecular weight of DNA as 660 g/mol/base pair. Based on recommendation from Ellison et al. (10), limit of detection (LOD) and quantification (LOQ) were established for each assay; LOD as the lowest concentration of the standard dilutions returning at least one positive replicate out of the three replicates prepared, LOQ as the lowest concentration at which all three positive replicates were able to amplify on the purified target dsDNA (Fig. S2). Efficiency of all qPCRs with standards was 97-103%. In the field study, when one or more qPCR replicate amplified, the average cycle threshold (C_t_) value of all replicates showing amplification is reported (Table S3).

All positive samples from the tank experiment 1 (microcosm), tank experiment 2 (true eDNA), and the field collections showed a similar sigmoidal PCR amplification curve. A subset of the positive samples was sequenced in order to confirm that the sigmoidal PCR amplification curve represented the target species. Species authenticity of *S. mansoni* was confirmed by amplicon sequencing in 22% of all the positive samples in the tank experiment 1, in 12.5% of the positive samples in the tank experiment 2, and in 80% of all the positive field samples. This was done by purifying qPCR products using Qiagen MinElute PCR purification kit (Qiagen, USA), followed by cloning using Topo TA cloning kit (Invitrogen), as described by Sigsgaard et al. (11), and finally sequencing of the inserted PCR fragment (Macrogen, Europe). All DNA extraction blanks and PCR controls performed throughout this study were negative.

